# Phasor based single-molecule localization microscopy in 3D (pSMLM-3D): an algorithm for MHz localization rates using standard CPUs

**DOI:** 10.1101/191957

**Authors:** Koen J.A. Martens, Arjen N. Bader, Sander Baas, Bernd Rieger, Johannes Hohlbein

**Affiliations:** Laboratory of Biophysics, Wageningen University and Research, Stippeneng 4, 6708 WE Wageningen, The Netherlands; Laboratory of Bionanotechnology, Wageningen University and Research, Bornse Weilanden 9, 6708 WG Wageningen, The Netherlands; Microspectroscopy Research Facility, Wageningen University and Research, Stippeneng 4, 6708 WE Wageningen, The Netherlands; Faculty of Applied Sciences, Delft University of Technology, Lorentzweg 1, Delft, 2628 CJ Delft, The Netherlands

## Abstract

We present a fast and model-free 2D and 3D single-molecule localization algorithm that allows more than 3 million localizations per second on a standard multi-core CPU with localization accuracies in line with the most accurate algorithms currently available. Our algorithm converts the region of interest around a point spread function (PSF) to two phase vectors (phasors) by calculating the first Fourier coefficients in both x- and y-direction. The angles of these phasors are used to localize the center of the single fluorescent emitter, and the ratio of the magnitudes of the two phasors is a measure for astigmatism, which can be used to obtain depth information (z-direction). Our approach can be used both as a stand-alone algorithm for maximizing localization speed and as a first estimator for more time consuming iterative algorithms.

## Introduction

Single-molecule localization microscopy (SMLM) has become a widely used technique in the biomolecular sciences since seminal contributions successfully demonstrated a roughly ten-fold improvement in spatial resolution over conventional fluorescence microscopy^1–3^. The key concept of SMLM is that the position of a single fluorescent emitter can be determined with an accuracy exceeding the diffraction limit as long as the emission of different molecules is sufficiently separated in time and space^4–6^. To localize the individual particles with sub-diffraction accuracy in two or three dimensions, a number of approaches have been developed^7^. Frequently employed localization algorithms involve the use of two-dimensional Gaussian functions to fit the intensity profile of individual emitters with high precision. These approaches, however, tend to be slow due to their iterative nature^8,9^, albeit data analysis in real time using graphics processing units (GPU) has been successfully demonstrated^10^. Faster localization algorithms using, for example, center of mass (CoM) calculations^11^ or radial symmetry^12,13^ tend to have lower localization accuracy or lack the ability to assess 3D information at > 10^5^ localisations per second^14^. Although a Fourier domain localization scheme for non-iterative 2D localization has been demonstrated theoretically, that method has not been widely adopted as it did not offer significant improvements in either localization speed or accuracy compared to iterative algorithms^15^.

Here, we introduce a simple and non-iterative localization algorithm with minimal computation time and high localization accuracy for both 2D and 3D SMLM. Our approach is based on the phasor approach for spectral imaging^16^. In pSMLM-3D, we calculate the location and astigmatism of two-dimensional point spread functions (PSF) of emitters. The real and imaginary parts of the first coefficients in the horizontal and vertical direction of the discrete Fourier transformation represent coordinates of the *x*- and *y*-phasors in a phasor plot. The associated angles provide information on the *x*- and *y*-position while the ratio of their magnitudes is a measure for astigmatism that can be used to determine the z-position of the emitter after introducing a cylindrical lens in the detection pathway of the microscope^17,18^. Our analysis of simulated PSFs with different photon counts indicates that phasor-based localization achieves localization rates in the MHz range, using only the CPU rather than requiring a GPU implementation, with similar localization accuracy as Gaussian-based iterative methods. Next to this, we localized microtubules in dendritic cells in three dimensions obtaining similar results with pSMLM-3D as with an iterative Gaussian-based algorithm. Finally, we implemented our algorithm both as a stand-alone MATLAB script and into the freely available ImageJ^19^ plug-in ThunderSTORM^20^ to which we further added the possibility to calculate intensity and background levels of emitters based on aperture photometry^21^.

## Methods

Data analysis in SMLM consists of the following steps: Identifying potential molecules and selecting regions of interests (ROIs) around their approximate localization, sub-pixel localization within the ROI, and visualization of results (Fig. 1a). Here, we will only focus on the sub-pixel localization step. We simulated the intensity pattern of a point source emitter using a full vectorial model of the PSF as described previously^22^ and depict it pixelated and with shot noise, mimicking a typical camera acquisition under experimental conditions (Fig. 1b). As our algorithm is able to utilize astigmatism commonly introduced by placing a cylindrical lens in the emission path for localization in three dimensions^17,18^, we simulated the full-width at half-maximum (FWHM) of the PSF in *y*-direction to be larger than in *x*-direction. We then calculated the first Fourier coefficients in the *x*- and *y*- direction by isolating them from the full two-dimensional discrete Fourier transformation of the ROI (see also SI: S1). Although the coefficients can also be calculated without calculating the complete Fourier transformation, this did not improve localization speed in the MATLAB environment. The real and imaginary part of each first Fourier coefficient are the coordinates of a phasor, which both are fully described by their phase angles (Θ_*x*_ and Θ_*y*_) and magnitudes (*r_x_* and *r_y_*), representing the relative position of the emitter in real space and values for the PSF ellipticity, respectively (Fig. 1c). To aid the reader, we calculated the inverse Fourier transformation using only the isolated first Fourier coefficients to show the data which is used for calculation of the emitters position and widths in real-space (Fig. 1d). We also show the localized position as determined from the phasor plot (Fig. 1d, green cross) and the ground-truth position (Fig. 1d, pink cross). The two elements represented in the phasor plot (Fig. 1c) have different distances to the origin. These magnitudes are inversely proportional to the FWHM of the original PSF: *r_y_* < *r_x_*, leading to FWHM_*y*_ > FWHM_*x*_, in agreement with the simulated data. The ratio of the PSF width in *x*- and *y*-direction can be used to calculate unknown *z*-positions of emitters in sample data after recording of calibration data.

**Figure 1:**
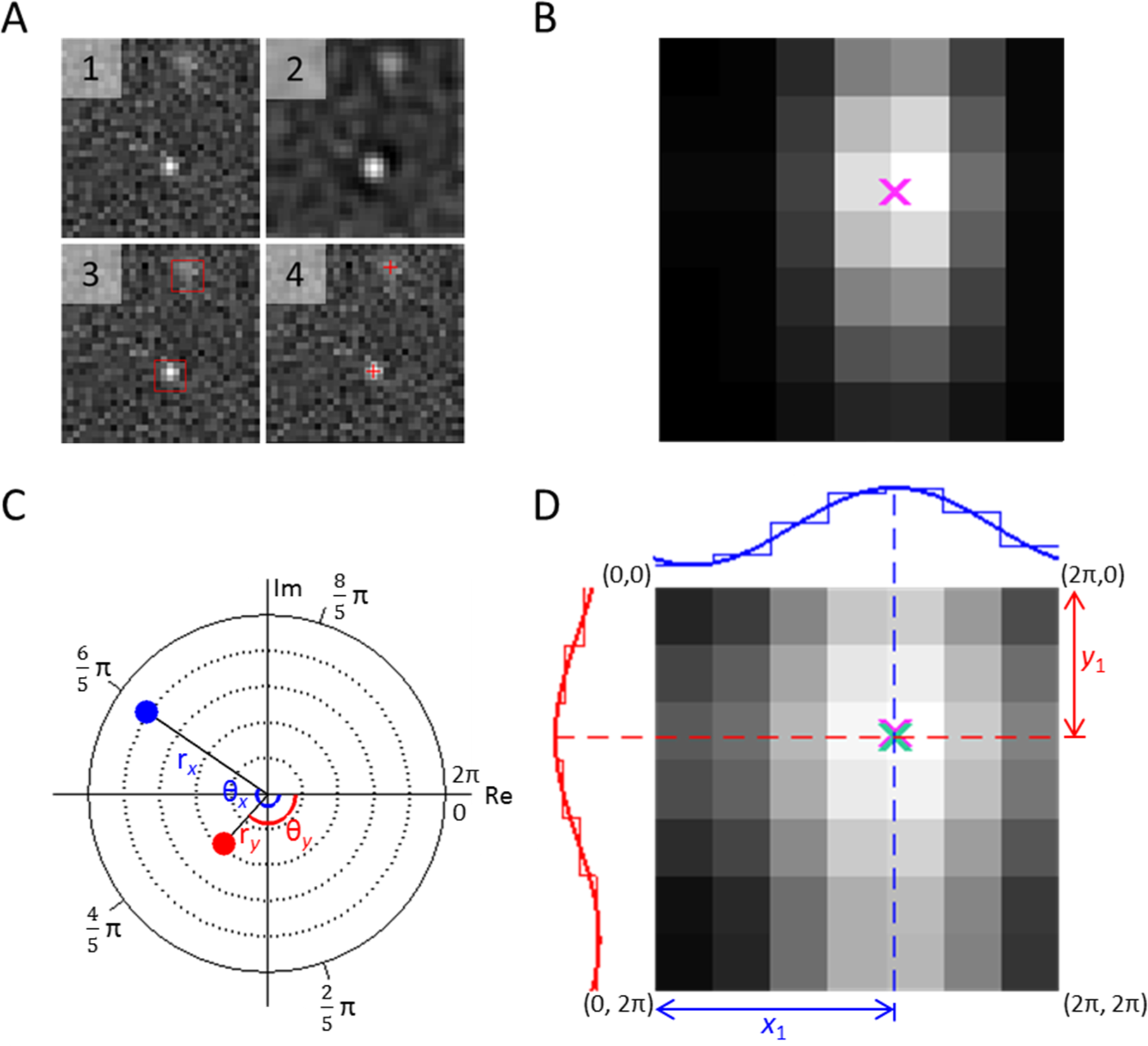
Illustration of sub-pixel localization using the phasor approach. **(A)** Standard workflow in single-molecule localization microscopy: (1) Acquisition of raw image data; (2) Image filtering; (3) Approximate localization of emitters: obtaining ROIs; (4) Sub-pixel localization. **(B)** Strongly pixelated image (7×7 pixel) including noise representing standard conditions using camera-based detection of a simulated ellipsoidal point spread function with the ground-truth localization indicated by a pink cross. **(C)** Phasor plot representation of the two first Fourier coefficients of the image data. By plotting their real versus the imaginary part, the angles Θ_*x*_ and Θ_*y*_ represent the position (phase) of the molecule in real image space as the markings on the straight circle in the Fourier domain indicate the normalized 1D position of the true center. Furthermore, the magnitudes *r_x_* and *r_y_* are reciprocally related to the PSF width in *x* and *y* in real space, respectively. Dotted lines are added for visual guidance. **(D)** Inverse Fourier transformation of the two first Fourier coefficients with the cumulated discrete intensity profile plotted in *x*- and *y*-direction and fitted with a sinusoid for visual guidance. From the angles Θ_*x*_ and Θ_*y*_ obtained from (C) and plotted in (D), we obtain the position of the molecule in the image domain using *y*_1_and *x*_1_marked by a green cross, with the pink cross from the ground-truth position shown for comparison.

## Results

To assess the performance of the phasor algorithm, we analyzed simulated data with a background level of 10 photons/pixels and a varying degree of total photon counts from the emitter ranging from 80 to 50,000 photons using images of 15×15 pixels. We compared the localization speed and accuracy of pSMLM-3D with other well established localization algorithms (for details see SI: S2): Gaussian-maximum likelihood estimation (Gauss-MLE)^10^, Gaussian-least squares fit (Gauss-LS)^23^, radial symmetry^12^ (RS) and centroid^11,23^ (Fig. 2). We further included the Cramer-Rao Lower Bound (CRLB) to indicate the theoretically achievable resolution where relevant^24^.

**Figure 2:**
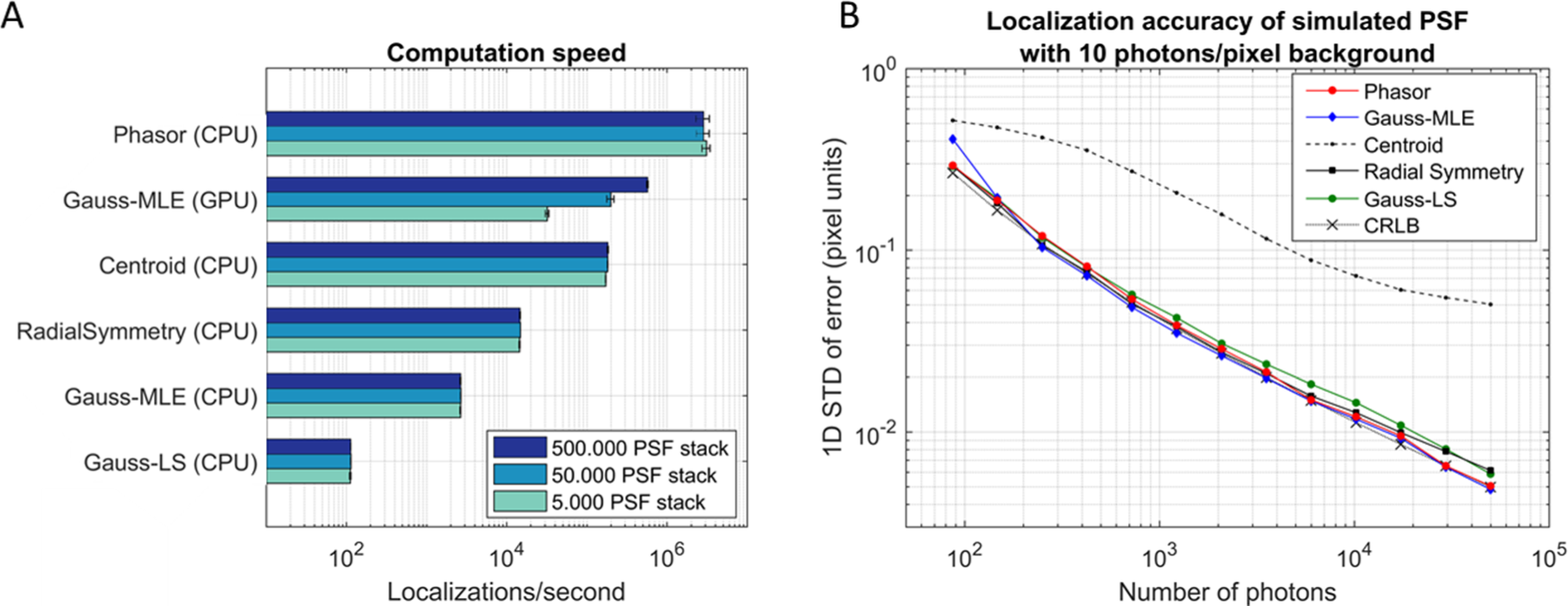
Comparison of computation speed and localization accuracy of phasor with other localization algorithms (Gaussian-MLE^10^, Gaussian-LS^23^, radial symmetry^12^ and centroid^11,23^).**(A)** Speed of localization after loading the raw data in the memory in MATLAB. 7×7 pixel ROIs are used; the amount of PSFs at once supplied to the method is varied. **(B)** Accuracy comparison of phasor localization with other localization algorithms, comparing simulated PSFs with different total photon counts on a 10 photon/pixel background. Accuracy in the horizontal direction of all methods together with the Cramer-Rao lower bound^24^ are shown. ROI size is 5×5 (<10^3^ photons) or 7×7 (>10^3^ photons) pixels for the phasor algorithm, and 7x7 pixels for all other algorithms.

In terms of localization speed, pSMLM-3D achieved more than 3 million localizations per second (3 MHz) when using ROIs with 7×7 pixels (Fig. 2a). This localization rate is at least an order of magnitude faster than our adapted implementations of other CPU-based algorithms and even significantly faster than GPU-enabled Gauss-MLE. Moreover, we found that the localization speed of GPU-based algorithms depends on the amount of data transferred to the GPU: Whereas a stack of 5,000 7×7 pixel images was analyzed at a rate of 30 kHz, a stack of 500,000 images (representing 49 MB of transferred data to the GPU), could be analyzed at 600 kHz. For CPU-based algorithms, this dependency is absent, allowing fast analysis of small PSF-containing image stacks, indicative that CPU-based methods are well suited for real-time analysis.

To assess the localization accuracy of the different localization algorithms, we cropped the area around each simulated PSF (15×15 pixels) to create ROIs of 7×7 pixels (in line with the ‘rule of thumb’ fitting region size of 2 · 3*σ_PSF_* + 1)^10^ for analysis by all methods, except for phasor where we used ROIs of either 5×5 (for simulated photon counts < 10^3^ photons) or 7×7 (> 10^3^ photons). We note that determining the optimal ROI size is often challenging for all localization algorithms: albeit working with larger ROIs can potentially increase the localization accuracy as more information from the PSF is extracted, larger contributions from background and near-by other emitters can have a diametric effect. Moreover, these effects depend on the photon count of the PSF and the type of localization algorithm used for analysis (see also SI: S3).

The comparison showed that for PSFs consisting of 80 photon counts, the localization accuracy is around 0.3 unit pixels for Gauss-MLE, Gauss-LS, RS and phasor and reduces to 0.005 unit pixels at 50,000 photon counts in line with the theoretically expected improvement of the localization accuracy being proportional to the square root of the photon number^25^ (Fig. 2b, SI:S8 - Fig. S10A). Between these outer limits, pSMLM-3D shows on average a small 3.7% decrease in accuracy compared to Gauss-MLE. We further note that the computationally inexpensive centroid based localization algorithm has a substantially worse localization accuracy, in line with earlier results^12^. We repeated all simulations at reduced background levels of 1 or 5 photons per pixel showing that the localization accuracies of all methods improve with lower background levels (SI: S4).

So far, we limited our analysis to localizations in two dimensions. As our algorithm allows using the ratio of the relative widths of the PSF in *x*- and *y*-direction introduced by astigmatism, the position of an emitter in three dimensions can be determined after performing a calibration routine in which photostable fluorescent emitters (e.g. latex beads) are imaged at different focus positions. Compared to non-astigmatic PSFs, we used larger ROIs (11×11 pixels for phasor, and 13×13 pixels for other methods, see SI: S5 – Fig. S4 for details) to account for the larger PSF footprint. Comparison of phasor with other algorithms on simulated astigmatic PSFs showed that phasor remained the fastest tested algorithm whilst providing a lateral localization accuracy close to that of Gauss-MLE, and better than Gauss-LS and Centroid (SI: S5 – Fig. S5, SI: S8 – Fig. S10B). Although RS is capable of determining the ellipticity of PSFs^14^, localisation rates did not exceed 10^5^ Hz and the localisation accuracy did not match that of Gauss-MLE.

With Gaussian-based methods, the PSF FWHM can be elucidated directly from the Gaussian fit; in our algorithm, the phasor magnitudes depend not only on the PSF FWHM in the respective directions, but also on the background. This dependency can introduce a bias if the background of the calibration series differs from that of the actual data. However, the ratio of the phasor magnitude in *x* versus in *y* remained unaltered (SI: S6), indicating that calibration of the ratio between the magnitudes versus z-depth should be performed. We calculated the axial localization accuracies using phasor and Gauss-MLE both of which provide similar accuracies decreasing from around 200 nm at very low photon counts (<500 per PSF) to under 20 nm at high photon counts (>10,000 per PSF) (SI: S7, SI:S8 – Fig. S10C).

To demonstrate the effectiveness of pSMLM-3D, we performed a standard 3D-STORM measurement of fixed immature dendritic cells with fluorescently labeled microtubules. In total, we recorded 50,000 frames (256×256 pixels), resulting in 6.1 GB of raw data containing roughly 2.7 million localized molecules. We analyzed this data with the ThunderSTORM^20^ plugin for ImageJ^19^ both with phasor and Gaussian-MLE. Fig. 3A shows an overview of the localization data as acquired via ThunderSTORM-phasor and ThunderSTORM-Gauss-MLE. During image acquisition, the limited signal-to-noise ratio required changing the size of the ROIs for phasor and Gauss-MLE to 7×7 and 11×11 pixels, respectively. The lateral (Fig. 3B) and axial (Fig. 3C) resolving power of phasor is in line with that of Gauss-MLE. The complete analysis time using multi-core computing, including the filtering of the image to find potential single molecules and excluding the loading of the data in the computer’s memory, was over 5 hours for Gauss-MLE, while it took only around 90 seconds for pSMLM-3D. Entirely omitting sub-pixel localization shortened the computation time by only ~5 s, which means that around 95% of the 90 second computation time is spend on image filtering and obtaining the approximate localization. Complete SMLM analysis with phasor under these conditions is at over 500 frames per second, indicative that it is fast enough for real-time analysis applications (Fig. 3D).

**Figure 3:**
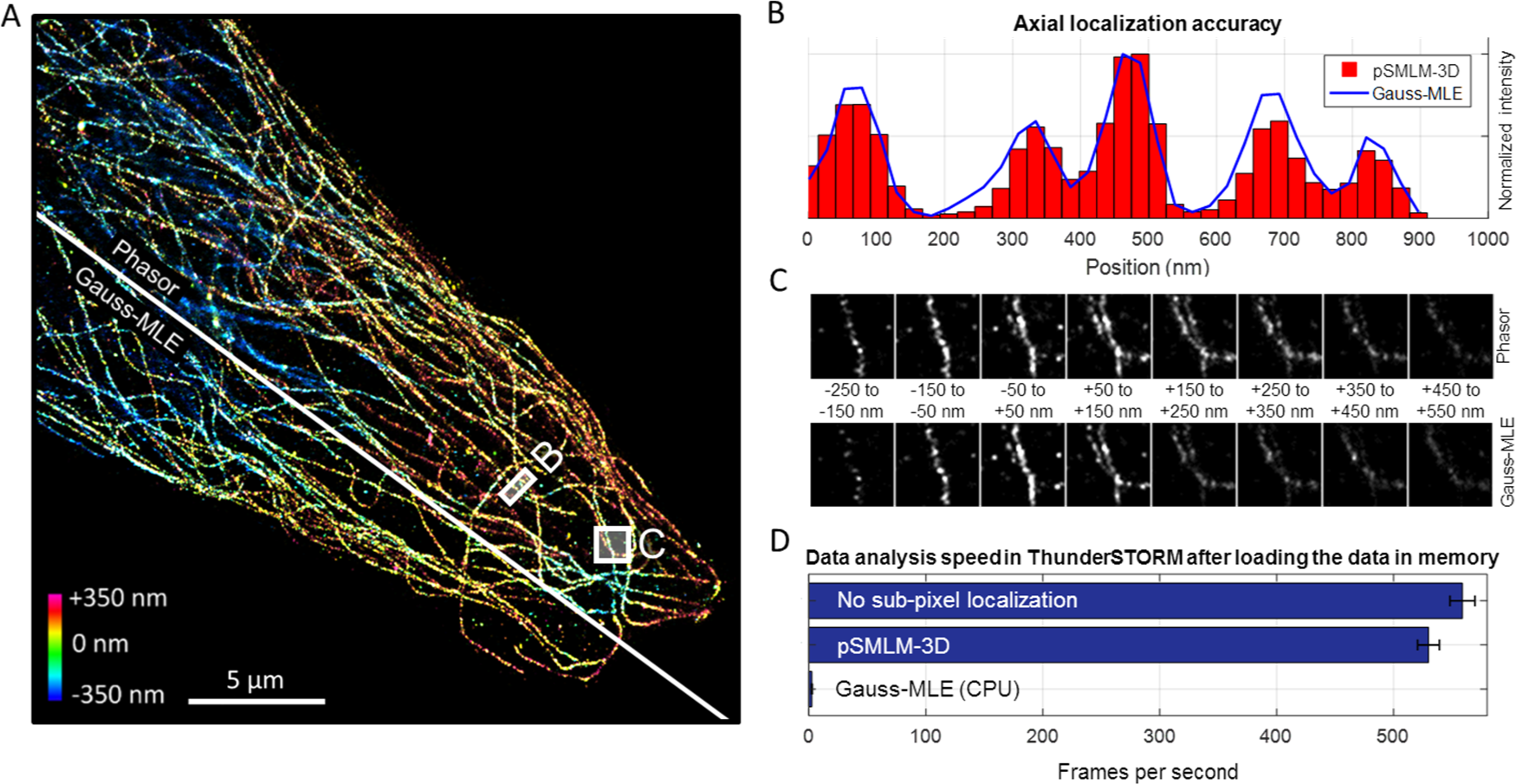
Analysis of a superresolved microtubule network of fixed HELA cells. **(A)** Visualization of superresolution data after ThunderSTORM analysis using phasor (top, 7×7 pixel ROI) or Gauss-MLE (bottom, 11×11 pixel ROI) as sub-pixel localization algorithm. Axial position is color-coded between -350 nm and +350 nm. Note that this does not encompass all localized fluorophores. **(B)** Lateral resolving power of phasor (red bars) and Gaussian-MLE (blue line). Shown here are three microtubule spaced below the diffraction limit taken from panel B in (A). **(C)** Axial resolving power of phasor (top) and Gaussian-MLE (bottom). Each subpanel shows localized fluorophores in a 100 nm window. **(D)** Localization speed of complete analysis (image filtering, approximate localization, and sub-pixel localization) using ThunderSTORM without sub-pixel localization (top), ThunderSTORM-Phasor (middle) and ThunderSTORM-Gauss-MLE (bottom). Error bars represent standard deviations of at least three repeats.

## Discussion and Conclusion

The presented pSMLM-3D combines excellent localization accuracies in three dimensions with exceptional localization speeds achievable on standard PCs. In depth analysis of synthetic point spread functions with different photon counts and background levels indicated that pSMLM-3D achieves a localization accuracy matching that of Gaussian-based maximum likelihood estimation even at low signal-to-noise ratios. Moreover, we demonstrated localization rates above 3 MHz, which is at least an order of magnitude increase in speed compared to other CPU-based algorithms. In fact, even compared to GPU implementations of Gaussian-based localization algorithms^26^, our algorithm is faster thus significantly reducing the computational barrier and costs to analyze experimental SMLM data. Porting the phasor approach to a GPU environment is likely to achieve only marginal improvements in speed as the bandwidth of transferring raw data is becoming a limiting factor. However, implementations using field programmable gate arrays (FPGAs) directly connected to the camera chip are feasible, with real-time SMLM analysis with Gaussian methods shown before^27^.

Subpixel localization rates in the MHz range satisfy even the most demanding applications as frame rates of cameras suitable for single-molecule detection are currently not above 100 Hz (full frame), indicating that phasor localization could be used in real-time environments. Moreover, some iterative localization algorithms currently use the centroid-based localization as a first estimation^10^. We believe that in that setting, the phasor approach can replace the initial step as it shows a speed as well as an accuracy improvement. We note that all necessary functions for performing the phasor algorithm are trivial, which allows for an easy upgrade of existing SMLM software packages. In computational environments in which a fast Fourier transformation function is not inherently present, a minimal algorithm to compute only the first Fourier coefficient can be written to minimize computation times, as we did for our JAVA implementation of phasor (SI: S9).

Compared to MLE, subpixel localization is possible in smaller areas around each emitter with good localization accuracy, allowing to use effectively a higher concentration of fluorescently active emitters. This is especially apparent with astigmatism, where a 11x11 pixel size in the phasor approach gives similar localization accuracy as 13x13 pixel size in the Gauss-MLE approach. This directly results in a possible increase of 40% in fluorophore density with the same chance of having partial emitter overlap. However, we note that our current phasor implementation does not provide means of resolving molecules whose emission partially overlap.

Like most localization algorithms currently available, pSMLM-3D assumes well-behaved PSFs with symmetrical emission profiles. Therefore, the algorithm depends on emitters having sufficient rotational mobility as emission profiles deviating from symmetrical PSFs can result in significant localization errors as has been discussed^28–31^.

In summary, we believe that pSMLM-3D holds great promise to replace or complement commonly used localization algorithms, as the combination of high localization speeds and high localization accuracy has not been shown to this extent before.

## MATERIAL AND METHODS

### PSF simulations

PSF simulations have been performed as described earlier^22^ with NA = 1.25, emission light at 500 nm, 100 nm/pixel camera acquisition and image sizes set to 15×15 pixels. The centre of the PSF is within ±1 pixel of the centre of the image, and in case of simulated astigmatic PSFs, the astigmatism has a FWHM ratio between 0.33 and 3.0. We used a full vectorial model of the PSF needed to describe the high NA case typically used in fluorescent super-resolution imaging. We accounted for the fact that in fluorescent super-resolution imaging the emitter can rotate freely during the excited state lifetime (~ns), so for many excitation-emission cycles an average over randomly distributed emission dipole orientations will be observed in one camera frame (~ms).

### Computer and software specifications

All computational work was performed on a 64-bit Windows 7 computer with an Intel Core i7 6800K CPU @ 3.40GHz (6 cores, 12 threads), NVIDIA GTX1060 GPU (1280 CUDA cores, 8 GHz memory speed, 6GB GDDR5 frame buffer, driver version 376.51), and 64 GB of DDR4 RAM on a ASUSTeK X99-E WS motherboard.

We used two software packages in this work: MATLAB (MathWorks, UK) version 2016b and FIJI^32^. FIJI is based on ImageJ^19^ version 1.51 n, using JAVA version 1.8.0_66.

### Software scripts used

Unless specified otherwise, we used variants of the phasor script we have written in MATLAB (SI: S9). JAVA-implementation of the phasor approach is based around a written minimal discrete Fourier transformation (SI: S10), and includes a aperture photometry-based method to estimate PSF intensity and background (also see SI: S11)^21^. Gauss-MLE, Gauss-RS, radial symmetry, center-of-mass, and Cremer-Rao lower bound algorithms were adapted from earlier uses^10,12^. For Gauss-MLE, 15 iterations were used^10^; Gauss-LS had 400 maximum iterations, with a tolerance of 10^−6^.

### Chemicals

All chemicals were purchased from Sigma-Aldrich and used without further purification, unless specified differently.

### Labeling of *in vivo* microtubules

Microtubule were fluorescently labeled via a double antibody labeling; primary antibody was a mouse-anti-βTubulin, clone E7, isotype mouse IgG1; the secondary antibody was labeled with Alexa 647 (Goat anti-Mouse IgG (H+L) Superclonal Secondary Antibody, Alexa Fluor 647, ThermoFischer).

The labeling procedure was as follows: HELA cells cultured on glass coverslips were fixed for 5 minutes with methanol at -20 °C, followed by 25 minutes fixation by 4% paraformaldehyde (PFA) in PBS. Next, a blocking step to prevent unspecific adsorption was performed by adding 3% bovine serum albumine in PBS pH 7.2 + 20 mM glycine (MP Biomedicals) and incubated for 1 h. Primary antibody was added and incubated for 1h. After washing with PBS, the secondary antibody was added and incubated for 45min. After a final washing step, the cells were post-fixed with 2% PFA in PBS for 15 min at RT and stored in PBS with 0.05% NaN3, with the final cells being stable for imaging for several days in PBS.

During imaging, a Gloxy buffer^33^ with added 35 mM β-mercaptoethanol, was added to boost blinking of the fluorophores. This blinking buffer was freshly prepared on the day of imaging.

### Single-molecule microscopy

We used a home-built microscope for imaging similar to a microscope described in more detail elsewhere^34^. Briefly, our microscope is equipped with a laser engine (Omicron, Germany), a 100x oil immersion SR/HP objective with NA = 1.49 (Nikon, Japan), and an Zyla 4.2 plus sCMOS camera for image acquisition (Andor, UK). 2x2 binning was used during acquisition, which resulted in a pixel size of 128x128 nm. A cylindrical lens with 1000 mm focal distance was placed in the emission path at 51 mm from the camera chip to enable astigmatic measurements; alignment of the lens’ optical axis was performed to ensure PSF elongation in *x*- or *y*-direction.

### Microtubule imaging and analysis

Fully labeled cells with added blinking buffer were imaged for 50.000 frames (256×256 pixels) at 10 ms frame time. A 642 nm laser at 70 mW in Hilo was used for imaging of the fluorophores, a 405 nm laser at increasing power throughout the measurement was used to activate fluorophores. Analysis was performed via the ThunderSTORM^20^ plugin for ImageJ^19^, with phasor added as sub-pixel localization option (SI: S10). ThunderSTORM parameters for image filtering and approximate localization were kept constant for phasor and Gauss-MLE localization: a β-spline wavelet filter with order 3 and scale 3 was used, and approximate localization was done via an 8-neighbourhood connected local maximum, with a peak intensity threshold equal to the standard deviation of F1 of the wavelet filter. These settings are the default ThunderSTORM settings; the only difference was a β-spline wavelet filter scale of 2 rather than 3.

Sub-pixel localization was performed with either elliptical Gauss-MLE (11×11 pixels, 1.6px initial sigma) or phasor (7×7 pixels). Localizations for pSMLM-3D and Gauss-MLE in the acquired datasets were filtered as follows: intensity/background > 2; background standard deviation < offset/2 (note that these are raw sCMOS counts rather than photon numbers). Calibration files were recorded under similar circumstances with immobilized fluorescent latex beads (560 nm emission, 50 nm diameter), and moving the piezo z-stage from -1000 nm to +1000 nm. These calibration files were used during the sub-pixel localization to calculate the z-position of the fluorophores.

Visualization of the superresolution data was done via the average shifted histogram options, with a magnification of 3 (Fig. 3A,C) or 5 (Fig. 3B). No lateral or axial shifts were added. 3D was enabled and visualized colored, after which a composite image was formed in FIJI (Fig. 3A).

## SUPPORTING INFORMATION

Supporting information is available online. For the latest implementation of pSMLM-3D into the ThunderSTORM plugin including documentation, please see https://github.com/kjamartens/thunderstorm/tree/phasor-intensity-1 and the specific folder “Compiled plugin”.

## ACKNOWLEDGEMENTS

J.H. acknowledges support from a Marie Curie Career Integration Grant (#630992). K.M. is funded by a VLAG PhD fellowship awarded to J.H. B.R. acknowledges support from European Research Council grant no. 648580. We thank Ben Joosten and Gert-Jan Bakker from Radboud UMC Nijmegen for kindly providing us with fixed dendritic cells and relevant labeled antibodies. We thank Sjoerd Stallinga for providing PSF simulation code. We thank Paul Reynolds and Christophe Leterrier for helpful discussions.

